# A Workflow for High Through-Put, High Precision Livestock Diagnostic Screening of Locomotor Kinematics

**DOI:** 10.1101/2022.02.04.479126

**Authors:** Falk Mielke, Chris Van Ginneken, Peter Aerts

## Abstract

Locomotor kinematics have been challenging inputs for automated diagnostic screening of livestock. Locomotion is a highly variable behavior, and influenced by subject characteristics (e.g. body mass, size, age, disease). We assemble a set of methods from different scientific disciplines, composing an automatic, high through-put workflow which can disentangle behavioral complexity and generate precise individual indicators of non-normal behavior for application in diagnostics and research. For this study, piglets (*Sus domesticus*) were filmed from lateral perspective during their first ten hours of life, an age at which maturation is quick and body mass and size have major consequences for survival. We then apply deep learning methods for point digitization, calculate joint angle profiles, and apply information-preserving transformations to retrieve a multivariate kinematic data set. We train probabilistic models to infer subject characteristics from kinematics. Model accuracy is validated for strides from piglets of normal birth weight (i.e. the category it was trained on), but the models infer the body mass and size of low birth weight piglets (which were left out of training, out-of-sample inference) to be “normal”. The age of some (but not all) low birth weight individuals is underestimated, indicating developmental delay. Such individuals could be identified automatically, inspected, and treated accordingly. This workflow has potential for automatic, precise screening in livestock management.

## 2 Introduction

Veterinary diagnostics have struggled with a methodological trade-off between high precision and high through-put. In the era of genomics, proteomics, and the like, the strive for accurate diagnostics of livestock diseases has directed considerable attention to the development of modern laboratory tests (Howson et al., 2017; Lamy and Mau, 2012). Conventional imaging techniques also play a role, but usually require special equipment and measurement techniques (e.g. radiography, microscopy, ultrasound, *cf*. Yitbarek and Dagnaw, 2022). These methods are high precision tools, but low through-put or expensive, some potentially invasive, and therefore not generally suitable for broad monitoring of farm animals. On the other hand, computational techniques are increasingly available to mine extensive data sets collected with sensors or cameras for diagnostically relevant signals (Gómez et al., 2021; Neethirajan, 2020; Netukova et al., 2021; Piñeiro et al., 2019; Wurtz et al., 2019). “Precision Lifestock Farming”, an application of integrated management systems, might be the desired economic model. These techniques complement the high precision tools by enabling broad screening and early detection of abnormalities, often preceding manual, veterinary intervention. Precision Lifestock Farming is promising in terms of its impact on animal welfare and economic success, but pitfalls remain (Azarpajouh et al., 2020; Wathes et al., 2008). The term “precision” might be misleading. In an animal management context, it refers to the availability of individual animal data, and the reduction of inefficient and thereby non-sustainable management. However, in practice, the use of sensors and cameras often still is restricted to superficial measures such as the overall activity or the mere occurrence or frequency of certain behaviors of individuals. For example, in swine farming, conventional video cameras can be used to monitor activity, and reduction can be associated with disease (Benjamin and Yik, 2019; Fernández-Carrión et al., 2017; Vranken and Berckmans, 2017); specificity and precision of these methods deserve further validation.

One class of behaviors that is typically monitored with such cameras is locomotion. Locomotion involves multiple subsystems, and one of the major challenges is to understand how exactly locomotor patterns are altered by conditions of the animal or by external circumstances. The involved subsystems are the musculoskeletal apparatus, energy supply, metabolism, and multiple levels of neuro-motor control. The kinematic and dynamic measurements obtainable by cameras and measurement equipment represent the collective output of interacting variables of the ensemble of subsystems (Nishikawa et al.,2007). In normal function, all of them are potentially affected in different, non-trivial ways by characteristics of the animal (Young and Shapiro, 2018), e.g. age (due to individual development), weight (due to body segment inertia), and size and morphology (due to allometrics in general and specific muscle lever relations in particular). In non-normal conditions or disease, another dimension of complexity is added. In consequence, studying alterations in specific locomotor patterns holds more diagnostic potential than activity measurement alone. Kinematic measurements have enabled the inference of many aspects of the locomotion of domestic animals (e.g. Netukova et al., 2021; Qiao et al.,2021; Schlageter-Tello et al., 2014; Serra Bragança et al., 2018); even individual recognition is possible in well-studied domestic species (e.g. Figueiredo et al., 2018; Patua et al., 2021). However, the cross-influence of the more or less correlated systems and co-variates mentioned above, and thus the superimposed effects of multiple factors, complicate data analysis and diagnostics. Most studies have relied on derived measures, such as speed or duty factor, as performance indicators, which neglects most of the individual movements of the joints and their temporal orchestration. For precision diagnostics, it would be desirable to have an automated system which tracks the locomotion of an individual, extracts and quantifies kinematics in all available detail, takes into account possible co-factors (such as age, size, and external physical conditions), and compares these observations to a reference for the species. Implicitly, this is what “a medieval husbandman”, i.e. a human classifier, would do with “his house cow or sow” (Wathes et al., 2008).

At the technical core of diagnostics is thus a classification problem: finding a diseased subset in a population of observations. Correct classification is complicated when there are multiple influence factors, but even more when the observation is subject to substantial intrinsic variability ^1^. Variability is a central feature of motor behavior: even for identical external conditions and in a single individual, it can be noted that “successive movements […] never exactly repeat themselves” (Bernstein, 1935). Could a putatively abnormal or pathological behavior actually fall “within the bell curve” of normal variability? How likely is that? Which of the many “input factors” is responsible, and how, for a given (temporary) alteration in the collective output? These analysis questions are common in research on bipeds (e.g. Bruton et al.,2013; Ganley and Powers, 2005; Stiffler-Joachim et al., 2020) and quadrupeds (e.g. Irschick and Jayne,1999; Pike and Alexander, 2002; Stavrakakis et al., 2014), and the solution is not novel. Multivariate models are capable of handling complex situations, given sufficient data. Multivariate *probabilistic* models (see below) are suited to also capture intra-individual variability and yield effect likelihoods. However, the high dimensionality of kinematic data sets, the multi-parameter, multi-level (hierarchical) covariate situations, and the high digitization workload have often been a limiting factor for the generation of quantitative models of vertebrate locomotion (Jackson et al., 2016; Michelini et al., 2020; Seethapathi et al., 2019).

Several recent technological advances have enabled researchers to tackle scientific questions on locomotion in a more efficient way. Firstly, the past few years have brought huge leaps in terms of computer vision, deep learning, and thereby semi-automatic video digitization methods (Corcoran et al., 2021; Jackson et al., 2016; Karashchuk et al., 2021; Mathis et al., 2020; Mielke et al., 2020). These tools typically require a manually digitized subset of the data as the “training set” for a neural network, which is then able to digitize further videos in high through-put, hopefully with reasonable accuracy. A second field of technological advance are the aforementioned probabilistic models, which build on an elegant computational implementation of Bayesian theory (Markov Chain Monte Carlo / MCMC sampling, *cf*.Gelman et al., 2020; McElreath, 2018; van de Schoot et al., 2021). Such models can naturally incorporate hierarchical parameter interrelations and intrinsic variability. The main reason for this is that probabilistic models work on data distributions, and their outcome are distributions and “effect likelihoods”, rather than point estimates. This can be informative on an intrinsically varying process such as locomotion (Mielke et al., 2018). Machine Learning methods for video digitization are validly advancing to be the standard in kinematic analysis, whereas probabilistic models still lack recognition in the field, despite their potential. To summarize, the mentioned advances in computer vision and statistical modeling enable us to (1) acquire a lot of quantitative data with minimal to no workload, and (2) model them in a suitable way. It would be desirable to adapt those technological advances for veterinary use, generating a classifier which could identify systematic alterations in the locomotion of domestic animals, and thereby enabling the computer-supported diagnostic screening for deficiencies, pathological states, and diseases.

Domestic pigs are a well-studied model system in which scientific interest joins the economic interest of commercial breeding. These animals have been subject to a variety of locomotor studies, including paradigms to test the effects of breed (Mirkiani et al., 2021), birth weight (Vanden Hole et al., 2018a,2021, 2017), surface friction (von Wachenfelt et al., 2008), welfare (Guesgen and Bench, 2017), various pathologies (Abell et al., 2014; Benasson et al., 2020; LaVallee et al., 2020), and more (*cf*. Netukova et al.,2021). Of particular interest has been the occurrence of a subset of individuals which are born with lower weight (LBW, low birth weight) than their “normal” (NBW) littermates. There are multiple standards to classify these birth weight categories, using absolute mass, litter quantile criteria, or asymmetry of body proportions (Amdi et al., 2013; D’Inca et al., 2011; Feldpausch et al., 2019; Quiniou et al., 2002; Roehe and Kalm, 2000; Van Tichelen et al., 2021; Wang et al., 2016). A possible cause of low birth weight is intra-uterine growth restriction, and LBW phenotype seems often, but not always, to correlate with low vitality and a reduced chance of survival (Baxter et al., 2008; Hales et al., 2013; Muns et al., 2013; Van Ginneken et al., 2022). Locomotor maturation after birth is quick (Andersen et al., 2016; Vanden Hole et al., 2017), yet crushing by the sow constitutes one of the major causes of piglet mortality (Edwards and Baxter, 2015; Marchant et al., 2000). The likelihood of being crushed is directly reduced by more agile locomotion. Thus, locomotor capabilities are crucial for piglet survival, and delayed development might be fatal.

Previous studies from our group (Vanden Hole et al., 2021, 2017) raised the hypothesis that the apparent difference in LBW and NBW individuals can be attributed to delayed development. They measured spatiotemporal gait variables (e.g. stride frequency and distance, speed, duty factor), which are collective variables of the actual kinematics (*cf*. Aerts et al., 2000; Newell and Liu, 2021; Nishikawa et al., 2007). This strategy has the advantage that it requires only five landmarks (four limbs, one reference) to be digitized, which used to be a crucial trade-off to handle large data sets. However, the the collective variables cannot capture full information on intra-limb coordination (i.e. the relative timing of segmental movements within a limb; as opposed to inter-limb coordination, i.e. the relative timing of the cyling of the different limbs). This complicates disentangling effects such as those of size, age, (birth) weight, and disease. It is expected that animals adapt their gait to the physical constraints of motor behavior, which are depending on the weight and other characteristics of the subject. However, the changes to kinematics might be more subtle, and collective variables might not be altered in a distinct way. For example, an animal might learn to move its joint angles in a more efficient way by adapting clearance to substrate conditions (von Wachenfelt et al., 2008), which could in principle be achieved without changing the speed of voluntary locomotion on those substrates. Hence, targeting authomated gait analysis and diagnostic classification of swine, it would be desirable to include full kinematic information.

Using the semi-automatic, machine-learning digitization techniques mentioned above, one can extend the analysis of gait variables to quantities of intra-limb coordination with manageable workload. However, using the whole set of raw point coordinates of joint points of interest raises the issue of dimensionality (two to three coordinates per reference point, simply too many data variables). Statistical modeling requires a minimum number of observations for being able to infer effects of the different variables (Austin and Steyerberg, 2015; Frick, 1996; Maxwell et al., 2017; Riley et al., 2020). The common solution is to reduce the dimensionality with an appropriate transformation. To choose a transformation, it can be exploited that common analysis procedures in locomotor biomechanics require steady state locomotion. “Steady state” implies that the behavior consists of repetitive blocks of kinematics, i.e. stride cycles. And one of the most common sets of techniques in physics and engineering to handle cyclic data is Fourier Analysis, or more specifically Fourier Series Decomposition (FSD; Bracewell, 2000; Fourier, 1822; Gray and Goodman, 1995; Mielke et al., 2019; Pike and Alexander, 2002; Webb and Sparrow, 2007). With FSD, joint angle profiles are transformend into their representation in the frequency domain, i.e. an array of harmonics. Some of the characteristics of the profiles (namely mean angle, amplitude, and phase) are more readily captured by those harmonics and can optionally be removed. This is most intuitive in the case of phase: removing phase differences enables a mathematically optimal temporal alignment of the profiles. By isolating the other characteristics, mean and amplitude, the joint angle profiles can be transformed to meaningful quantities such as dynamic posture (mean joint angle and effective range of motion), and coordination *sensu stricto* (relative phase/joint timing and residual kinematics, *cf*. Mielke et al., 2019). Harmonics are independent of temporal sampling and duration: the coefficient array is of fixed size, which is useful for subsequent multivariate analysis methods, such as Principal Component Analysis (PCA). Another advantage of this transformation procedure is that it is reversible because all mathematical information is retained in the process (which is not the case when using collective variables alone). This means that joint angle profiles can be reconstructed for any observed or hypothetical point in parameter space, which enables in-sample and out-of-sample predictive sampling.

To summarize, the Fourier Series decomposition provides a mathematically convenient and biomechanically meaningful representation of the kinematic data, which opens up new options for data analysis and modeling.

In this study, we establish a workflow which can be automated and used to identify individual animals locomoting differently from the “normal” reference, based on video recordings, deep learning digitization, mathematical transformations, and probabilistic modeling. A conventional, 2D kinematics data set is extracted with the aid of deep learning tools from lateral videos of walking piglets. By applying multivariate analysis and FSD, we separate spatiotemporal gait variables, dynamic posture, and coordination, and model their relation to subject characteristics (mass, size, age, and birth weight category). Crucially, this constitutes the complete information captured by locomotor kinematics, and all parameters are submitted to an inclusive, probabilistic model. As a test case, we tackle the question of whether low birth weight in domestic piglets is an indication of delayed development, and attempt to quantify the delay with an inverse modeling strategy as follows. Intuitively, and conventionally, joint kinematics are considered the output of the locomotor system. Therefore, conventional statistical models might consider them on the “outcome” side; on the “input” side, the effects of birth weight, age, speed, or other parameters are quantified. Herein, we use a different approach, and invert the model. We construct a probabilistic computer model which describes “age” and other subject characteristics as a function of all available kinematic parameters. The rationale is similar to that in subject recognition tasks: given a certain kinematic profile, can we infer (characteristics of) the subject? We split our data set into birth weight classes (LBW, NBW), and train the model on only the strides from NBW observations. This NBW model is our “kinematic reference” model, quantitatively capturing the expectation of what would be “normal” by inferring the plausible age range for a given kinematic observation. We then use that trained model to compute out-of-sample inference of individual LBW observations.

Our hypothesis is that, if LBW were at the same stage of postnatal locomotor development as their NBW siblings, then the model should accurately infer the age of the LBW animals. Conversely, if the LBW piglets are delayed in development, the model would underestimate their age. Thus, by applying this inverse modeling strategy and comparing the computer-inferred age to the actual age of the LBW piglets, we can quantify and potentially falsify a hypothesized delay in locomotor development.

The components of this classification workflow are not novel, and commonly used in physics and engineering. We use available machine learning tools to digitize videos, apply a series of well-known transformations, and train a probabilistic model classifier. We demonstrate that a set of individual locomotor events can be used to distinguish individuals which develop slower than expected, in a temporal accuracy of four to eight hours (which is a considerable timespan for neonate animals). These are precise diagnostic measurements, generated at high through-put, with the overall aim of improving animal welfare, all of which is in line with the prototypical ideal of Precision Livestock Farming.

## 3 Materials And Methods

### 3.1 Data Acquisition

Recordings were done at a local farm in Belgium during several trips in October and November 2017. Farrowing was monitored to select Topigs x PIC piglets for another experiment (Ayuso et al., 2021). Piglets from selected litters were weighed at birth and numbered with non-toxic skin markers. Low birth weight (LBW) was classified by birth weight quantile (lowest 10%of each litter) and by a maximum mass of 800g (D’Inca et al., 2011; Litten et al., 2003; Van Tichelen et al., 2021; Wang et al., 2016); all other piglets are assigned the NBW category. At variable time points afterwards (ages 1 – 10 hours), piglets were briefly taken from their pen and brought to a separate room for video recording (see below). Animals were recorded in pairs (as in Mielke et al., 2018), which drastically reduced anxiety and increased their motivation to cooperate. A few animals were recorded repeatedly, usually with a changing partner. Animals were ear-tagged and followed up: recording was repeated at approximately 4 and 10 days of age. That data was part of the digitization procedure (i.e. “deeplabcut” network training), but excluded from further analysis (i.e. probabilistic modeling, see below). The subject characteristics documented for analysis are birth weight (continuous, and categories “LBW”/“NBW”), mass at recording, age at recording (i.e. hours since farrowing), sex, and size. The size of the animal was approximated by a Principal Component Analysis (PCA) of digitization landmark distances along all segments (“size PCA”, only first PC used, 93%of variability). Size and mass are expected to correlate, yet deviations would indicate animals of particularly slender or rotund habitus. All procedures followed ethical regulations and guidelines, and were approved by the Ethical Committee for Animal Testing of the University of Antwerp, Belgium (ECD 2015-26).

The recording room contained an elevated runway (150 × 50cm), covered with a rubber mat to increase friction, and visible through a transparent frontal shield. Color videos were recorded (camera model: GC-PX100BE, JVC, Japan) at a temporal sampling rate of 50 frames per second and a spatial resolution of 1920 × 1080 pixels (later cropped to 500 px height), from a distance at which the field of view would exactly capture the entire runway. A chess board at the back wall enabled spatial calibration. Video surveillance was permanent during the presence of the animals and stopped only in between recording sessions. Animals were able to move freely on the enclosed platform. To stimulate locomotion, the two animals were repeatedly placed on opposite ends of the runway. Gentle tickling on the back and grunting vocalization of the researcher were other successful strategies to induce targeted locomotion in the direction perpendicular to the camera axis. After recording sessions the piglets were returned to their litter and remained with the sow. The workflow herein involved handling of the animals as a consequence of the research setting. However, note that the procedure could easily be automated for continuous data collection by a suitable pen arrangement (Meijer et al., 2014; Netukova et al., 2021; Stavrakakis et al.,2014).

### 3.2 Digitization

We used the software DeepLabCut (DLC, Mathis et al., 2018) for digitization of all video material. In addition, a custom made point tracking software (Mielke et al., 2020) was used to generate a training set. In total, our dataset contained 180 videos (more than 11 hours, 169 animals) of video. Our goal was to prepare a general DLC network which is capable of automatically tracking piglets at multiple ages, and which can be shared and re-used for subsequent research questions. This is why the full data set was used for digitization and for the calculation of some derived measures (size PCA). However, the analysis focus of this study (see below) was only a subset of the data (i.e. the 58 animals of the youngest age class). The video processing workflow, applied to the full data set, was as follows. To get a balanced training set, one stride of each of the animals was selected, and the video was cut, cropped to runway height, and optionally mirrored horizontally so that movement would always be rightwards. All videos were concatenated and submitted to the DLC training set generation. DLC was set to select 2552 frames from these videos, which were tracked in an external software and re-imported for training (80%training fraction). Seventeen landmarks (i.e. points of interest or “key-points”; usually joint centers, fig. 1) were digitized, representing all body parts visible on the lateral perspective (head: snout, eye, ear; back line: withers, croup, tail base; forelimb: scapula, shoulder, elbow, wrist, metacarpal, forehoof; hindlimb: hip, knee, ankle, metatarsal, hindhoof). We selected a “resnet 152” network architecture and trained for 540, 672 iterations (16 days of computer workload). The network was then applied to digitize the continuous, full video recordings twice: once in default direction and once horizontally mirrored, because training set was always rightward movement.

**Figure 1:**
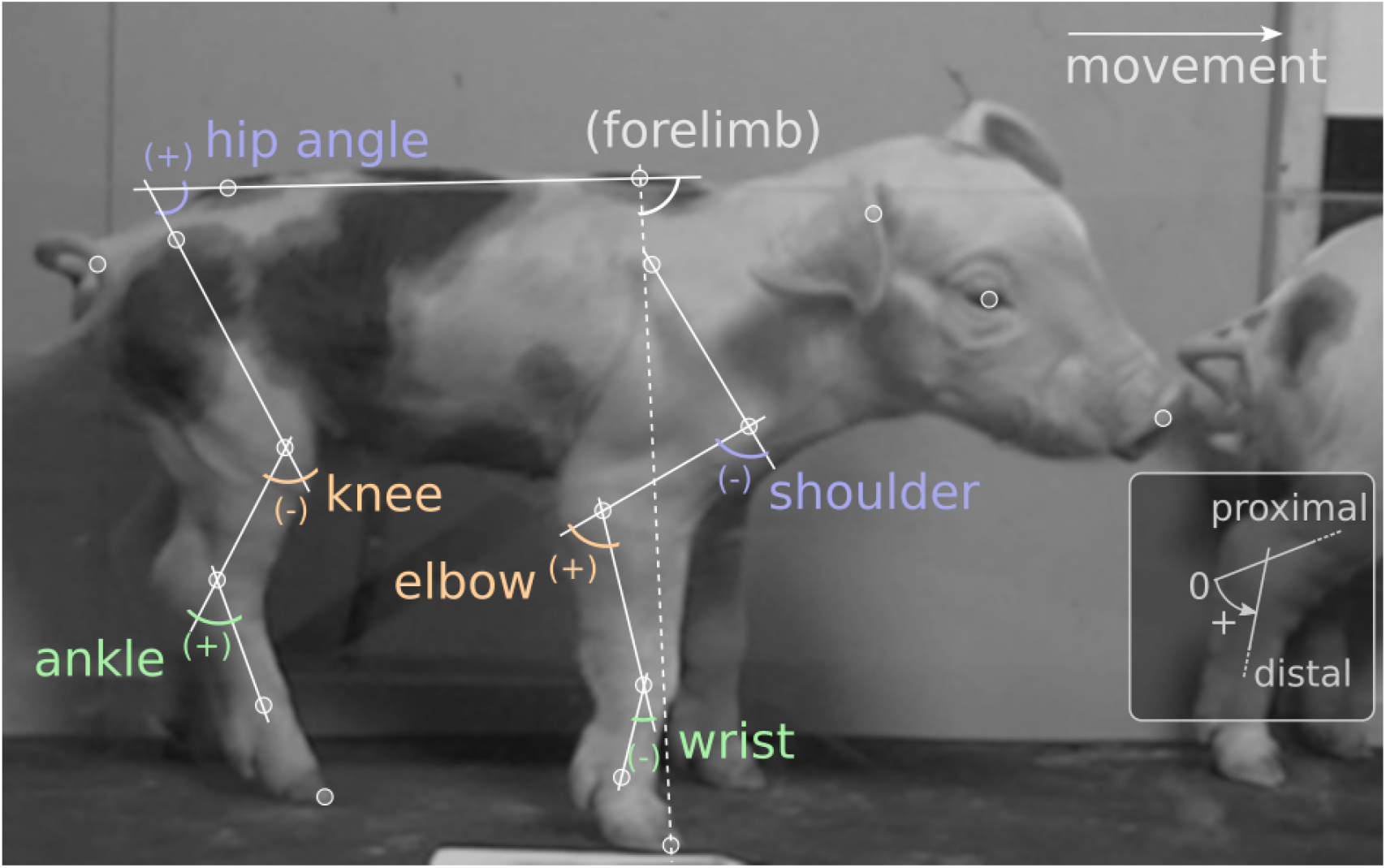
Video digitization and joint angle definitions. White circles mark points of interest (“landmarks”). Movement was always rightwards. Labels show joint angles, defined as shown in the inset: straight joint (parallel segments) corresponds to zero; counter-clockwise angles are positive. Forelimb angle was used as a reference for temporal alignment, but did not enter the analysis.

The next step is to find the relevant temporal sequences of walking in the continuous videos. Naturally, the trained network would only extract potentially useful landmark traces for episodes which resembled the training set, i.e. in episodes with a piglet moving perpendicular to the image axis, in lateral aspect and rightward direction. We automatically extracted 2597 of such sequences by filtering for high digitization “likelihood” provided by DLC, low noise (i.e. steady landmark movement) and consistent, plausible landmark distances. We further applied an automatic algorithm to find footfalls and label stride cycles in the candidate episodes (4730 cycles). This procedure involved a start-end-matching optimization (using Procrustes superimposition) to ensure that strides were indeed cyclical. To further assess digitization quality, gait variables were automatically extracted. Definition of these variables was chosen to simplify the automatic procedure, as follows. Stride distance, frequency, and speed are trivial measures of the animal movement. Duty factor is available for fore- and hindlimb, and measures the fraction of stride time in which the respective hoof is in ground contact. Clearance is approximated by quantifying the ratio of flexion of each limb (one minus the quotient of minimum and maximum absolute hip-toe-distance during the stride). Head and torso angle are the stride-average angles of the snout-ear or withers-croup lines with respect to the coordinate system. Hindlimb phase measures the time between hind- and forehoof touchdown, divided by the stride cycle duration. Where applicable, gait variables were prepared for analysis (see below) by converting them to dimensionless values (Alexander and Jayes, 1983; Hof, 1996) using the cumulated distance of landmarks along the snout-to-tailbase line of the animal as reference, extracted as stride average from the digitized landmarks. Only strides with plausible values (i.e. those which lie within the theoretical distribution of each parameter; 1862 cycles) where processed. Manual inspection further boiled down the data set to 897 stride cycles (the others excluded for digitization errors, multi-animal confusion, non-walking gait, intermittent or sidewards locomotion, or incompleteness).

Finally, 368 of the remaining strides from 58 animals were in the youngest age category (< 10 *h*) and thus selected for the present analysis, the data table is available online (see below).

### 3.3 Data Processing

The landmark data provided by DLC was further processed for analysis. Python code for the whole procedure is available (https://git.sr.ht/~falk/piglet_fcas, Python version 3.10.8 at time of model calculation, https://www.python.org). First, joint angle profiles (i.e. joint angle values over time) were extracted for all relevant joints and for the total forelimb angle (croup-withers-hoof). Shoulder, elbow, wrist, hip, knee, and ankle were the six joints sufficiently well digitized and therefore considered relevant for analysis. We then applied Fourier Series decomposition in the framework we previously termed Fourier Coefficient Affine Superimposition (FCAS, Mielke et al., 2019), a flexible procedure which subsumes the following steps. Joint angle profiles are cyclic, i.e. periodical, and can therefore be transformed to the frequency domain with a Fourier Series decomposition (8 harmonics were deemed sufficient by visual comparison of raw and transformed/retransformed profiles). In the frequency domain, the affine components (mean, amplitude, phase) of a joint angle profile are easily accessible (*cf*. Mielke et al., 2019). The forelimb angle served as reference to temporally align all cycles in the data set (removal of phase differences between different cycles; forelimb angle was not used further). Then, mean and amplitude of the joint oscillations were isolated for all joint angles and are categorized as “dynamic posture” parameters. Mean joint angle is the temporal average, whereas amplitude is related to effective range of motion (eROM). The residual, i.e. differences captured by non-affine Fourier coefficients, can be categorized as “coordination”*sensu stricto* (it measures the precise temporal succession of joint configurations). In our case, there were 96 variables of coordination (6 angles, 8 harmonics, real and imaginary) which were submitted to a PCA. Only the first 12 coordination components (*CC*) were used for statistical analysis, capturing 80.2%of the variability in coordination.

To summarize, FSD and FCAS served three purposes: (i) temporal alignment of the cyclic traces, (ii) separation of meaningful parameter categories (dynamic posture and coordination), and (iii) preparation for multivariate analysis via PCA. Basic script code (Python, Matlab and R) to perform FCAS can be found on a dedicated git repository (https://git.sr.ht/~falk/fcas_code).

Information retention is generally a strength of this method. FCAS and PCA are mathematical transformations, which means that the information content after transformation is theoretically identical to that prior to transformation (theoretically, because only a finite number of harmonics can be used, yet this is of little concern for continuous, smooth joint angle profiles). The neglected PCs and the residual not captured by 8 harmonics were the only information from kinematics of the given joints to be lost in this procedure, and by definition these contain the least information. Apart from that, all information present in the raw joint angle profiles enters the analysis. Though we used a 2D dataset herein, the procedure could be applied equally well to angles measured from 3D coordinate data (Scott et al., 2022).

Furthermore, all transformations are reversible, hence any analysis outcome can be translated back to kinematics with high accuracy. Reversibility bares a lot of herein unused potential, for example for interpolating unobserved subject states or for inferring kinematics of fossile species by phylogenetic and morphometric bracketing. Reversibility can also be of use when presenting raw joint angle profiles and their averages, as follows. One crucial aspect of the FCAS procedure is temporal alignment of the joint angle profiles in the frequency domain. In conventional temporal alignment, a single characteristic point in the stride cycle is chosen as a reference, wherein this is only “characteristic” for a certain part of one limb (e.g. left hindlimb hoof touchdown). Temporal alignment to the hindhoof touchdown might cause distinct peaks in the forelimb angle joint profiles to occur at different relative points in the stride cycle (e.g. ankle joint profiles in Fig. 3 below, lower half, green traces). If profiles show such variable peak positions, then their average will have a wider, less pronounced (i.e. lower amplitude), and potentially unnatural peak. For illustration, this is analogous to averaging two sine-waves of identical amplitude, but phase shifted: in the worst case, they cancel each other out (as in “destructive interference”). The problem is not restricted to pronounced peaks, but generally occurs if the temporal intra-limb coordination varies within a data set. Using FCAS, it is possible to get a more representative average of the raw traces which has its amplitude conserved, but phase and mean angle averaged. This is enabled by transformation to the frequency domain, separation of affine components, removal of phase differences by shifting to average phase, profile averaging, followed by inverse transformation back to the time domain. Because a set of profiles and phases may be calculated for each angle individually, and because phase relations can differ between joints, there are the options to align based on one reference angle (e.g. the whole forelimb, as done herein) or minimize all phase differences across all joints. Chosing the first option herein has implications: when plotting hindlimb joints aligned by a forelimb reference (as in Fig. 3, lower half), phases still differ, and the “destructive interference” problem might hamper averaging. In such cases it is possible to apply an extra, joint-wise FCAS alignment for the sole purpose of generating meaningful averages.

**Figure 2:**
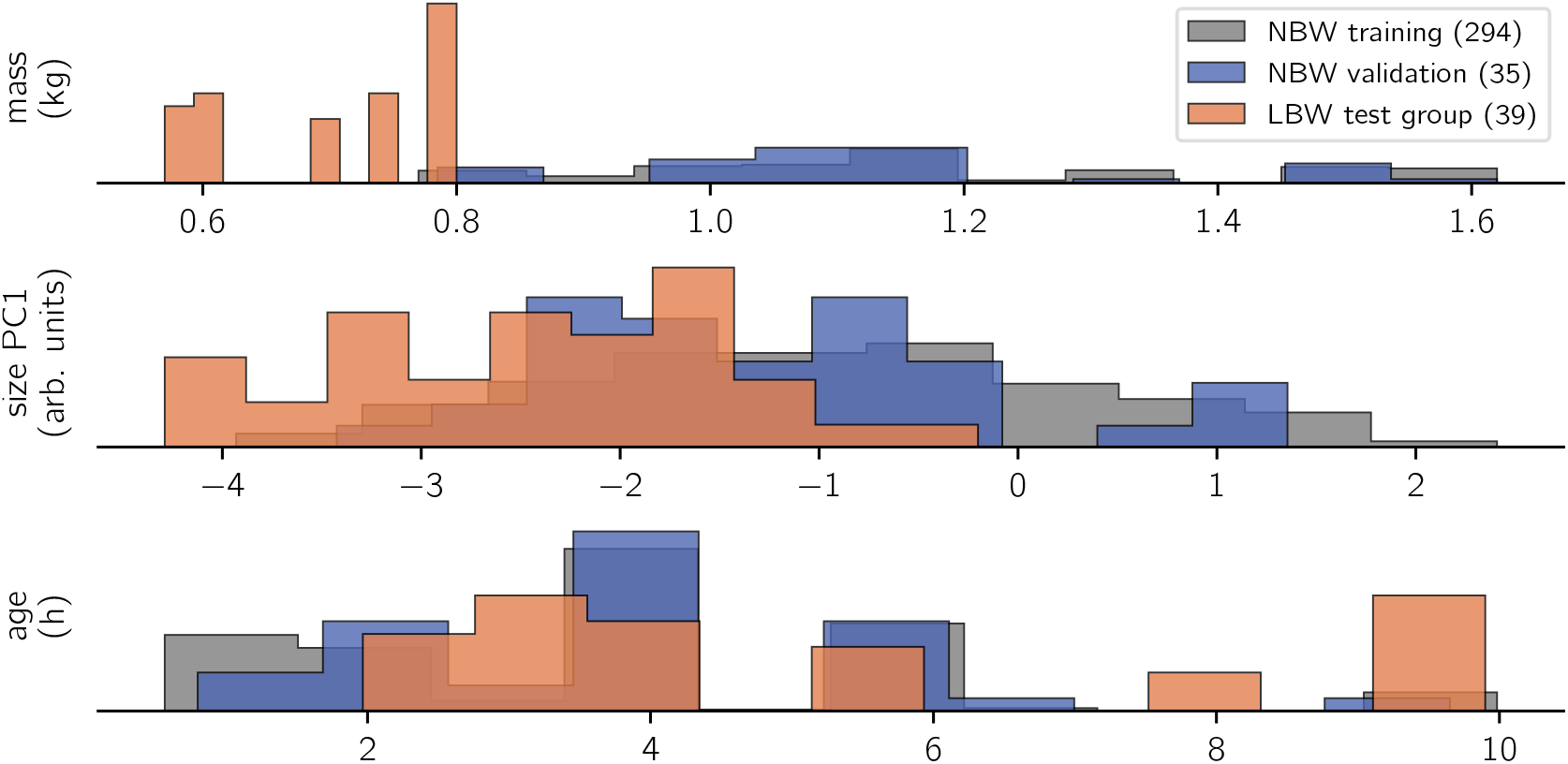
Histogram of observations. Trivially, the LBW group measured the lowest body masses in the data set. This correlated with a lower body size, whereas age is rather uniformly sampled for all study groups. Recordings happened opportunistically within the first ten life hours of the animals, repeated measurements were possible. Number of strides per class are indicated in brackets on the legend. Bar heights are scaled by sample size to show relative value distributions.

**Figure 3:**
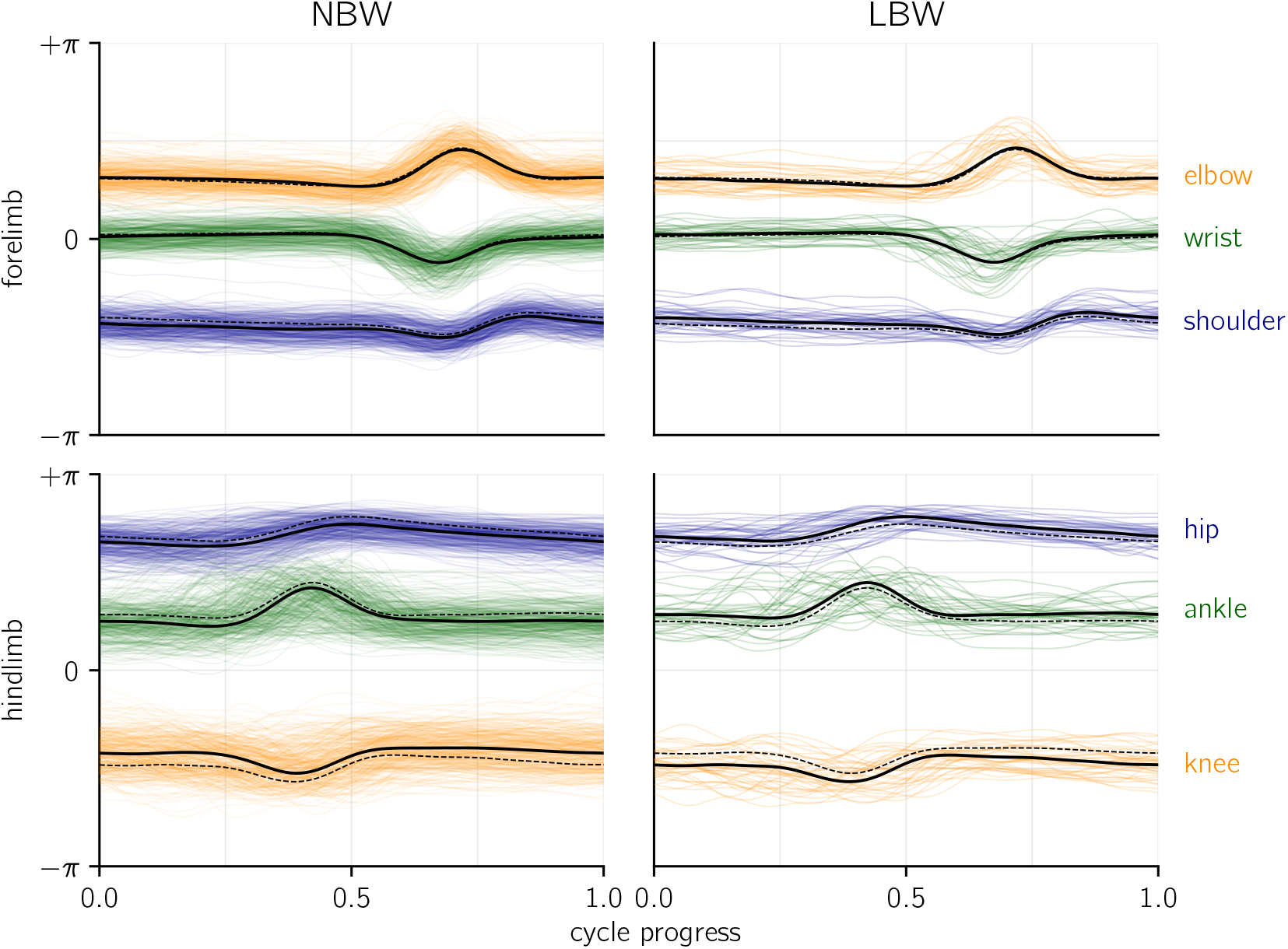
Joint angle profiles per joint, grouped by birth weight category. An angle of zero would be a fully extended (i.e. straight) joint. Thick lines represent the average profiles, dashed lines indicate the average of the opposite birth weight group for comparison. Colored, shaded lines show all raw profiles available for the present analysis. Temporal alignment was done based on total forelimb angle (see methods), yet for the shown hindlimb averages (but not for the raw profiles), a separate alignment of the hindlimb was performed.

### 3.4 Statistical Modeling

To summarize, four categories of variables were used for analysis:

- subject characteristics: age, sex, mass, birth weight category, size
- spatiotemporal gait variables: distance, frequency, speed, clearance (fore-/hindlimb), duty factor (fore-/hindlimb), head angle, hindlimb phase
- dynamic posture: mean joint angles and eROM for six joints
- coordination: the residual after extraction of dynamic posture (see above)

Our guiding question for model design is whether a probabilistic, linear model is able to infer subject characteristics (specifically: age, mass, and size) from raw kinematics (expressed as dynamic posture and coordination) and gait variables (collective variables). Given the common conception that kinematics are a complex output of an individual motor system, this might be considered an “inverse” modeling approach. The present analysis focused on three outcome variables (Fig. 2): mass (*kg*), size (*arb. units*, from a PCA of marker distances), and age (h). Though these outcome variables were specific per individual and recording session, we analyzed them “per stride” (i.e. there were multiple strides with identical subject measures on the outcome side).

The model formula is:

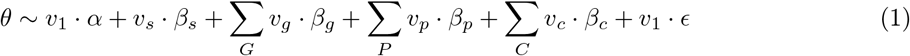

Herein, *θ* is either of the outcome subject characteristics, *β* are slopes associated with the model parameters (*s* sex, *G* gait variables, *P* dynamic posture, C coordination), *v* are data vectors (e.g. *v*_1_ is a vector of ones for the intercept *α* and model residual *03F5;*, and *v_s_* is a boolean vector coding for subjects of ‘sex == male‘). Priors (i.e. *a priori* assigned distributions) for all slopes were Normal distributions with mean and standard deviation corresponding to the mean and two times standard deviation of all observed values of each parameter; logarithmic transform was applied where necessary. The observable (“likelihood”) prior for *θ* was a Student’s T distribution (allows for wider-than-normal tails and robust regression) with a Gamma distributed *v* (degrees of freedom); *ϵ* was modeled to be a Half Cauchy distribution. The model was implemented using the Python library “PyMC” (version 4.2.2, Salvatier et al., 2016).

To re-emphasize, dynamic posture and coordination together effectively capture all the kinematic information of the stride. Hence, we train the predictor model with all kinematics, gait variables, and sex. Birth weight category (LBW, NBW) is a filter parameter: we split our data set into LBW strides and two NBW subsets (training and validation). Training is performed by MCMC sampling (‘sample‘ function in PyMC), and a No U-Turn sampler was set to sample with 32 chains, each 2^14^ tuning and equally many sampling steps. All post-hoc model checks confirmed convergence (inspection of traces, *bfmi* > 0.94 for all chains, Gelman-Rubin statistics ≈1 for all parameters, sufficient effective sample size). Model comparison was performed, iteratively leaving out model parameters or replacing some by meaningful combinations (e.g. duty factor combined for fore- and hindlimb). However, because we follow an “all in” strategy, the results have little instructive value for model construction: we might thus have retained parameters which are numerically unimportant for the NBW-only models.

The data set of *N* = 368 strides was split into three categories: (i) the NBW training set as reference with *N* = 294 strides, (ii) the NBW validation set (*N* = 35 strides), which is a random subset of NBW strides, approximately equal in size to (iii) the LBW test set with *N* = 39 strides.

The model was thus trained with a set of 294 NBW training strides (i). Inferences (model “predictions”) were then done per stride, for all observed strides (NBW training, NBW validation, and LBW test), iteratively using the ‘pymc.sample_posterior_predictive‘ function in PyMC after setting all the data arrays to the actual observed values for one given stride (using ‘pymc.set_data‘). The number of predictions usually matches the number of training samples, which means that all posterior information is used to construct the prediction distributions. We would thus retrieve mass, size, and age predictions (i.e. probabilistic inference) for each stride in the data set, which were then compared to the known, actual mass, size, and age.

All procedures, code, data, and this manuscript are available online (https://git.sr.ht/~falk/piglet_fcas).

## 4 Results

The present analysis is centered around a linear model which is designed to infer mass, size, and age (subject characteristics) from an extensive set of kinematic parameters from 2D videos. The numbers provided by the model sampling are equally extensive, and will only be reported in brief. The key purpose of the model is posterior predictive sampling of the LBW strides which were left out of the model, and which are analyzed in detail below.

To assess whether there are qualitative differences between the birth weight categories, one can compare the joint angle profiles (i.e. raw, angular kinematics) on which the present analysis was performed (Fig. 3). The intra-group variablility clearly exceeds the differences between groups, although it must be emphasized that groups are inhomogeneous (with regard to age, speed, etc.), which might lead to a bias if composition of LBW and NBW data differs. LBW walk with a more flexed hindlimb posture, as indicated by the parallelly offset average hip, knee, and ankle profiles. Additionally, NBW individuals on average seem to set the shoulder at a more extended angle. No differences in coordination are apparent (which would manifest in altered temporal structure of the profiles). These findings indicate that LBW kinematics are hardly distinguishable from NBW kinematics by qualitative, visual assessment, which is at least in part be due to high variability.

A quantitative comparison of variable kinematic measurements can be achieved with probabilistic linear models. For the purpose of predictive sampling (see below), we train models to describe the interrelations of kinematic parameters and subject characteristics in NBW piglets. The outcome of MCMC sampling of a linear model are value distributions for slopes, which in our case indicated how certain kinematic parameters are associated with a change in mass, size, and age (supplementary material 7.1). Of the gait- or coordination parameters, only hindlimb clearance was correlated with differences in animal mass. Mass was also associated with changes in the dynamic posture of the hip and ankle. For size, the model inferred associations with head angle, hindlimb duty factor and clearance, and one coordination component (CC3), as well as changes in the fore- and hindlimb posture and an effect of sex. Finally, age was associated with an increase in forelimb clearance, potential changes at the hip and wrist, and several coordination components (CC9, CC11). Some eROM slope distributions for age were high in average magnitude, but variable (the “credible interval” contained zero). These model results provide detailed insight into parameter interrelations in the present data set and indicate which of the parameters are the relevent ones to infer a given subject attribute in predictive sampling.

Performing in-sample and out-of-sample predictive inference with the models trained on NBW strides elucidated if and how left-out strides differed from NBW model expectation (Fig. 4). Note that, to capture variance (i.e. uncertainty in the prediction), each stride was sampled repeatedly.

**Figure 4:**
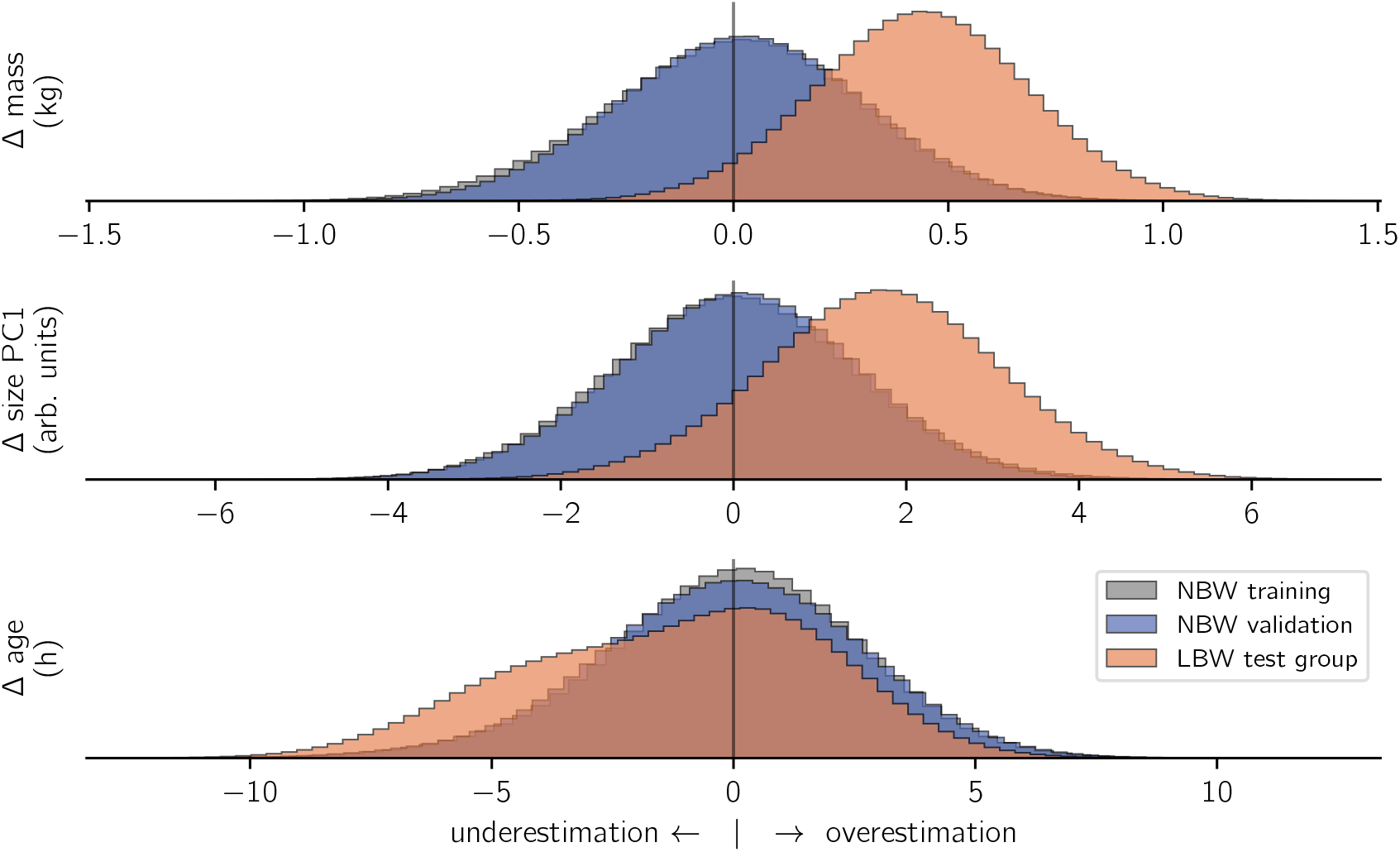
Model inference. For all included subject characteristics, models which were trained on NBW strides correctly inferred the training data (gray) and values from the validation set (blue). In contrast, the same models wrongly inferred the characteristics of LBW subjects (orange). The x-axes show the difference (Δ) between actual and predicted values per prediction. To facilitate comparison, histogram heights are again normalized per category.

Out-of-sample inferences for the *NBW validation set* matched those of in-sample NBW inference in terms of average values and standard deviation for all modeled outcome variables, which confirms that inference of subject characteristics from kinematics is possible. In contrast, inferences for *LBW strides* did not match those of the NBW training set. Low birth weight animals were inferred to be on average 0.44 kg heavier than actual, and their size was overestimated (+1.71 units). Both faults matched the actual differences in magnitude (*cf*. methods, Fig. 2). In contrast, the age inference for the low birth weight subjects were not normally distributed: most ages were correctly inferred from stride-wise kinematics, but ages for some strides were underestimated. The underestimation of those strides quantified to just below five hours.

In summary, the NBW-trained model “guesses” the size and mass of the animals producing LBW strides to be “normal” (although they are not), which indicates that these defining features of LBW do not reflect in altered kinematics. However, age inference is non-normal, i.e. some strides are classified as typical for animals of younger than actual age.

To find out whether the offset age inference was related to certain individuals, or certain strides from different individuals, we grouped the inferences per stride or subject and calculated the chance of over- or underestimating age. Of the 8 low birth weight subjects who contributed 39 strides, 4 individuals were consistently underestimated (Tab. 1). Consistently means that more than 75%of all predictive samples were below actual age, and that the ages for a majority of strides were on average underestimated. The magnitude of underestimation was between two and five hours. Curiously, those were the individuals recorded at a slightly higher age (> 5 hours). Overestimation in the other four LBW individuals was also consistent, but less so (less extreme underestimation rate, mean Δ < 2 h). Standard deviation of the estimates did not vary across individuals or birth weight categories.

**Table 1:** Age inference per LBW animal (compared to NBW average, last row). Δ: “inferred - actual” difference. Underestimation is defined as Δ < 0, “count”: per stride, “rate”: per predictive sample. h: hours, std: standard deviation.

| piglet | age<br>h | strides | underestimation<br>count | underestimation<br>ratio | pred. mean $\Delta$<br>h | pred. std<br>h |
| --- | --- | --- | --- | --- | --- | --- |
| b23 | 2.0 | 6 | 0 | 0.29 | 1.13 | 2.00 |
| b15 | 2.9 | 5 | 0 | 0.37 | 0.68 | 1.96 |
| b76 | 3.1 | 4 | 0 | 0.39 | 0.57 | 2.01 |
| b74 | 4.2 | 7 | 1 | 0.40 | 0.52 | 1.97 |
| 1794.5 | 5.6 | 5 | 5 | 0.90 | -2.57 | 1.99 |
| b58 | 7.8 | 3 | 3 | 0.91 | -2.85 | 2.00 |
| b19v2 | 9.8 | 1 | 1 | 1.00 | -6.14 | 1.99 |
| b56 | 9.9 | 8 | 8 | 0.99 | -4.58 | 1.96 |
| <all NBW> | <3.8> | 329 | 158 | 0.49 | -0.03 | 1.95 |

We conclude that underestimation of age is consistent over multiple strides of the same individual, and thus individual-specific.

## 5 Discussion

Quadruped terrestrial locomotion is the collective output of an ensemble of organismal subsystems, which is both reason and challenge for its usefulness in veterinary diagnostics. On one side, the kinematics can be quantified in multidimensional data sets, capturing the many degrees of freedom of the limb joints. On the other side, kinematic quantities are context dependent and affected by numerous subject characteristics (age, weight, pathologies,…) which also cross-influence each other. The challenge emerges to find the right trace of a given (or unknown) condition in the multidimensional observation on the background of kinematic variability. Deep Learning methods for video digitization have become available, and probabilistic computational models offer a flexible framework to mirror complex parameter relations. Once trained to a given question, these computer tools can achieve comparative diagnostic classification with minimal human interaction, e.g. for continuous screening in a farm setting. Multivariate systems have been a challenge to integrated management and precision farming, and the presented locomotor analysis workflow highlights a possible way to succeed in that challenge.

In this study, we have demonstrated a test case for generating a probabilistic model of piglet locomotion which incorporates all kinematic information.

Our example model was trained on a high number of observations which are considered “normal”, and applied to classify untrained observations in terms of deviation from normal behavior. The data stems from laterally filmed videos of normal (NBW) and low birth weight (LBW) piglet locomotor behavior from unrestricted walking gait (an inexpensive, high-throughput arrangement, and a common behavior). Low birth weight is often associated with low vitality (Baxter et al., 2008; Hales et al., 2013; Muns et al.,2013), and this supposedly correlates with deficient locomotion. Hence, the obvious first research question is whether birth weight has an influence on the locomotor behavior. Top-down, direct, visual assessment could justify the hypothesis that LBW walking kinematics are somehow different from “normal” (D’Eath,2012). Yet that is (i) hard to assess due to high behavioral variability and (ii) trivially expected given the adaptation to different physical properties of their body: gravitational force is a predominant constraint of locomotion, and it simply scales with animal weight. Our results showed that the eight LBW individuals we submitted to the weight-kinematics model were all over-estimated in terms of their weight, by the amount that matched LBW-NBW weight difference (Fig. 4). The same is true for the size model. This indicates that LBW, at least all those in our data set, are capable of walking as if they were of normal birth weight and size. This is the first example of a diagnostic model application: the model confirms quantitatively normal locomotor behavior despite occurrence of a given non-normal co-variate (weight).

A second diagnostic application is the identification of individuals (or even strides) which systematically deviate from an expectation or norm. Probabilistic models do not only classify “normal” or “not”: they yield a distribution of plausible values, and thereby a likelihood that a given observation is indicative of a problem. The same model architecture as above, but configured to infer age from a kinematic measurement, estimated some (but not all) individuals to be of lower than actual age (Tab. 1). Those were specifically the older of the LBW individuals, whereas the youngest ones (< 4h) walked as expected for neonates. Though we cannot fully rule out chance with our limited sample size, this provides evidence that the quick postnatal development was halted in those individuals. Our interpretation is that, at birth, LBW individuals putatively had the same capabilities as their NBW siblings, yet at least some “fell behind” regular development in the first hours. We can think of two possible reasons for this: (1) the birth process as a trauma might mask the actual capabilities of all neonates alike, concealing actual, pre-existing differences (Litten et al., 2003); (2) development is impeded by depleted energy reserves and a failure in (kin) competition and the perinatal struggle for teats and warmth (Le Dividich et al., 2017). We found little support for the first possible reason: top-down locomotor development is quick for both groups (Vanden Hole et al., 2018a, 2017), and muscular architecture shows no differences (Vanden Hole et al., 2018b). On the other hand, there is evidence for quick depletion of energy levels in the low birth weight individuals, which rectifies within a period of ten hours (Vanden Hole et al.,2019). This finding is consistent with the present study and supports the perinatal struggle hypothesis. Delayed development does not necessarily corroborate the hypothesis of locomotor deficiency in LBW. We would expect truly deficient strides to be substantially different from the data trained to the model, thus be either excluded or misclassified. Exclusion means that the used Deep Learning implementation could not capture deficient strides, or only in a way which led to exclusion in subsequent (automatic) quality checks (see below). We acknowledge that there currently still is room for refinement in the Deep Learning digitization procedure. Yet in the likely case that some deficient strides passed quality checks and were subjected to the model, we would expect them to be more “unpredictable” (i.e. higher variance of posterior samples). Instead, in our data set, inferences were consistent for repeated measures of an individual, without notable increase in variance across inferences per stride. For the affected subjects, we can even quantify a plausible delay of less than five hours, which could nevertheless be critical given the rapid maturation of locomotor behavior in this species (Vanden Hole et al., 2017) and the importance of postnatal competition. Such detailed information is valuable when evaluating the success of different mitigation strategies (e.g. supplementing energy to piglets, Schmitt et al., 2019). It must be emphasized that, just like other computational diagnostic tools, the method outlined herein is not intended for standalone use. Instead, it is complementary to or can facilitate the in-depth inspection. Nevertheless, the specificity of the presented gait analysis supersedes mere activity analysis: to our knowledge, being able to automatically retrieve an individual, probabilistic measure for developmental delay in swine has not been achieved before. Information retention is a feature of the presented workflow which we think can enable researchers and veterinaries to differentiate a multitude of potential influences on locomotor behavior, given sufficient reference data and an appropriate model design.

Our data set is limited and potentially biased in terms of LBW observations. There are much fewer valid LBW strides in our data set, in absolute numbers: only 39 of 368 observations are LBW. This could be interpreted as evidence for a lower capacity (despite equal potential) of LBW to produce normal locomotion. Yet there are proximal, trivial explanations: for this study, the 10%lower quantile of birth weights in a litter is considered LBW, and there is a hard cap of 800g. The resulting share is equal in our training set for video digitization, and in the final data set, because of pseudo-random, opportunistic sampling on-site (i.e. recording work was permanent, yet determined by farrowing and feeding of the subjects). The minority of LBW training videos might lead to an under-learning of those animals in the digitization network, which could lead to reduced digitization quality and therefore an exclusion bias for “non-normal” individuals. Though it seems unlikely, we cannot rule out reduced locomotor capacity in LBWs: the present data set is unsuited to count the occurrence of locomotor behavior due to its automatic generation. On the other hand, the strict stride filtering criteria for “good” kinematics may have involuntarily filtered out deficient individuals. Our conclusion that low birth weight individuals are non-deficient is strictly tied to the definition of the low birth weight category, which is herein based on weight criteria and did not regard phenotypical indicators of intra-uterine growth restriction (which we did not record, *cf*. Amdi et al., 2013).

A corollary question is which patterns in the kinematic variables cause the different age inferences. We report high magnitude (but also highly variable, i.e. “non-significant”) slopes inferred from the age model (supplementary material 7.1). Note that these slopes solely reflect effects within the NBW data subset. We also observed slight differences in the average hindlimb dynamic posture (Fig. 3). In fact, a more flexed hindlimb is typical for the youngest animals of both birth weight categories. We emphasized potential differences in group composition to explain that (e.g. sex effect in the “size” model), and different age per group might be a proximal explanation for the non-normal age inference in LBW. However, the average age of LBW animals (5.3 h) in our data set is nominally above that of NBW (3.8 h), which is a discrepancy with the age underestimation. Yet if we assume that the hypothesis of delayed locomotor development is correct, the nominal age would be misleading, and LBW effectively behave similar to younger animals. This can explain the apparent discrepancy in age group composition and age inferences from kinematics. It also suggests that dynamic posture might be the major proxy for perinatal maturation, though many other parameters also entered the probabilistic model and influenced the model outcome.

To summarize, we herein assembled state-of-the-art computer techniques for the purpose of individual diagnostics in quadruped locomotion, which we think constitute a valuable workflow for livestock screening and management. All components require some manual and computational efforts for initialization (network training, model regression). However, once that is done, the workflow is as follows:

- generate more video recordings (e.g. in an instrumented runway)
- apply the trained Deep Learning network for automated digitization
- identify stride cycles (automatic with framewise Procrustes comparison)
- stride cycle quality filtering by automatic criteria (end-start difference, constant speed,…)
- Fourier Series Decomposition, temporal alignment, and parameter transfromation (PCA)
- probabilistic classification (i.e. posterior predictive sampling) with an inverted model structure
- validation of above-threshold classifications

Except for the last (crucial) step, all of this can be fully automated, and the whole workflow is readily available for precision livestock farming. Monitoring can happen automatically (as in Litten et al., 2003;Netukova et al., 2021), which reduces delay in identifying individuals in need of intervention. Multiple models can be tested in parallel: in the present test case, the “weight” and “size” models found LBW locomotion indistinguishable from the “normal” reference group, whereas the “age” model specifically identified those animals which likely experience a delay in locomotor development. Likewise, tests for specific diseases could be set up. A more extensive (longitudinal) data set and more specific models are required to bring this tool into “clinical” or economical/commercial use, and one purpose of the present study was also to give sufficient explanations and references for readers unfamiliar with the mentioned methods. Nevertheless, we demonstrated that the modeling workflow is able to provide a high precision, high throughput method for domestic pig locomotor diagnostics.

## Supporting information

full kinematic raw data and meta data

## 6 Acknowledgements

The authors would like to thank Miriam Ayuso for organizing and participating in the recording sessions, as well as Laura Buyssens, Georgios Petrellis, Gunther Vrolix, Charlotte Vanden Hole, Denise Vogel and all students who joined for help during recordings. Maja Mielke provided valuable comments on the manuscript text.

## 7 Supplements

### 7.1 Supplements: Detailed Modeling Results

Asterisk (*) indicates slopes for which the credible interval did not include zero. FL: forelimb, HL: hindlimb, dyn.p.: dynamic posture, coord.: coordination, diml.: dimensionless, d.s.: dimensionless stride, eROM: effective range of motion

|  | category | parameter | age (h) | age (log) | size PC1 | mass (kg) |
| --- | --- | --- | --- | --- | --- | --- |
| 0 | model | intercept | +4.86 | +1.68 | −3.25 | +1.38 |
| 1 | subject | female → male | +0.05 | +0.00 | −0.38 * | −0.04 |
| 2 | gait | log. FL clearance | −0.10 | −0.08 | +0.09 | −0.02 |
| 3 | gait | log. HL clearance | +0.94 * | +0.32 * | −0.65 * | −0.15 * |
| 4 | gait | FL duty factor | +1.11 | +0.37 | +0.18 | +0.07 |
| 5 | gait | HL duty factor | +0.67 | +0.24 | +0.85 * | +0.11 |
| 6 | gait | d.s. distance | +1.32 | +0.31 | +0.45 | +0.18 |
| 7 | gait | d.s. frequency | +3.12 | +1.43 | +1.62 | +0.15 |
| 8 | gait | diml. speed | −1.10 | −0.64 | +0.82 | +0.00 |
| 9 | gait | hindlimb phase | −2.04 | −0.56 | −1.98 | +0.05 |
| 10 | gait | head angle | +0.26 | +0.14 | +1.44 * | +0.18 |
| 11 | dyn.p. | mean hip angle | +2.55 * | +0.59 * | −1.07 * | −0.45 * |
| 12 | dyn.p. | hip eROM | +2.64 | +0.94 | −1.34 | −0.52 * |
| 13 | dyn.p. | mean knee angle | +0.67 | +0.10 | −2.01 * | −0.12 |
| 14 | dyn.p. | knee eROM | −1.85 | −0.22 | −1.66 * | −0.13 |
| 15 | dyn.p. | mean ankle angle | −1.18 | −0.59 | +0.44 | +0.23 * |
| 16 | dyn.p. | ankle eROM | −1.92 | −0.88 * | +2.98 * | +0.60 * |
| 17 | dyn.p. | mean shoulder angle | +1.06 | +0.07 | +0.47 | +0.06 |
| 18 | dyn.p. | shoulder eROM | −0.34 | −0.17 | −0.65 | +0.01 |
| 19 | dyn.p. | mean elbow angle | −1.00 | −0.55 | +2.79 * | +0.20 |
| 20 | dyn.p. | elbow eROM | +3.22 | +0.96 | +0.11 | −0.30 |
| 21 | dyn.p. | mean wrist angle | −2.08 * | −0.98 * | +1.69 * | +0.04 |
| 22 | dyn.p. | wrist eROM | −0.24 | +0.21 | −0.84 | +0.16 |
| 23 | coord. | CC1 | +0.60 | +0.31 * | +0.10 | −0.05 |
| 24 | coord. | CC2 | +0.50 | +0.15 | −0.24 | −0.08 |
| 25 | coord. | CC3 | −0.25 | +0.00 | +0.88 * | +0.07 |
| 26 | coord. | CC4 | −0.62 | −0.26 | +0.13 | +0.04 |
| 27 | coord. | CC5 | −0.84 | −0.25 | −0.16 | +0.04 |
| 28 | coord. | CC6 | −1.49 | −0.51 * | −0.10 | +0.07 |
| 29 | coord. | CC7 | +0.24 | +0.22 | −0.76 | −0.08 |
| 30 | coord. | CC8 | −1.19 | −0.24 | −0.51 | +0.02 |
| 31 | coord. | CC9 | −2.96 * | −0.83 * | +0.66 | +0.06 |
| 32 | coord. | CC10 | −0.22 | +0.06 | −0.60 | +0.02 |
| 33 | coord. | CC11 | −2.05 * | −0.67 * | +0.10 | +0.08 |
| 34 | coord. | CC12 | −0.03 | −0.10 | +0.35 | −0.01 |
| 35 | model | $\epsilon$ | $\pm 1.81$ | $\pm 0.56$ | $\pm 0.98$ | $\pm 0.20$ |

## Footnotes

1 Whether “intrinsic” just describes variability for which no influence factor has yet been determined is a valid, but philosophical question beyond the scope of this study.

## References

Abell, C. E., Johnson, A. K., Karriker, L. A., Rothschild, M. F., Hoff, S. J., Sun, G., Fitzgerald, R., and Stalder, K. J. (2014). Using classification trees to detect induced sow lameness with a transient model. Animal, 8(6):1000–1009. doi: 10.1017/S1751731114000871.

Aerts, P., Van Damme, R., Van Elsacker, L., and Duchêne, V. (2000). Spatio-temporal gait characteristics of the hind-limb cycles during voluntary bipedal and quadrupedal walking in bonobos (Pan paniscus). American Journal of Physical Anthropology, 111(4):503–517. doi: 10.1002/(SICI)1096-8644(200004)111:4<503::AID-AJPA6>3.0.CO;2-J.

Alexander, R. M. and Jayes, A. S. (1983). A dynamic similarity hypothesis for the gaits of quadrupedal mammals. Journal of Zoology, 201(1):135–152. doi: 10.1111/j.1469-7998.1983.tb04266.x.

Amdi, C., Krogh, U., Flummer, C., Oksbjerg, N., Hansen, C. F., and Theil, P. K. (2013). Intrauterine growth restricted piglets defined by their head shape ingest insufficient amounts of colostrum. Journal of Animal Science, 91(12):5605–5613. doi: 10.2527/jas.2013-6824.

Andersen, A. D., Sangild, P. T., Munch, S. L., van der Beek, E. M., Renes, I. B., Ginneken, C. v., Greisen, G. O., and Thymann, T. (2016). Delayed growth, motor function and learning in preterm pigs during early postnatal life. American Journal of Physiology-Regulatory, Integrative and Comparative Physiology, 310(6):R481–R492. doi: 10.1152/ajpregu.00349.2015.

Austin, P. C. and Steyerberg, E. W. (2015). The number of subjects per variable required in linear regression analyses. Journal of Clinical Epidemiology, 68(6):627–636. doi: 10.1016/j.jclinepi.2014.12.014.

Ayuso, M., Irwin, R., Walsh, C., Van Cruchten, S., and Van Ginneken, C. (2021). Low birth weight female piglets show altered intestinal development, gene expression, and epigenetic changes at key developmental loci. The FASEB Journal, 35(4):e21522. doi: 10.1096/fj.202002587R.

Azarpajouh, S., Díaz, J. A. C., and Taheri, H. (2020). Precision livestock farming: automatic lameness detection in intensive livestock systems. CABI Reviews, 2020. doi: 10.1079/PAVSNNR202015031.

Baxter, E., Jarvis, S., D’Eath, R., Ross, D., Robson, S., Farish, M., Nevison, I., Lawrence, A., and Edwards, S. (2008). Investigating the behavioural and physiological indicators of neonatal survival in pigs. Theriogenology, 69(6):773–783. doi: 10.1016/j.theriogenology.2007.12.007.

Benasson, I., Wagnac, E., Diotalevi, L., Moore, D., Mac-Thiong, J.-M., and Petit, Y. (2020). Gait analysis of a post induced traumatic spinal cord injury porcine model. In 2020 42nd Annual International Conference of the IEEE Engineering in Medicine Biology Society (EMBC), pages 3803–3806.

Benjamin, M. and Yik, S. (2019). Precision Livestock Farming in Swine Welfare: A Review for Swine Practitioners. doi: 10.3390/ani9040133.

Bernstein, N. A. (1935). The Problem of the Interrelation of Co-Ordination and Localization. In Whiting, H., editor, Human Motor Actions: Bernstein Reassessed (1984), volume 17 of Advances in Psychology, chapter 2, pages 77–119. North-Holland.

Bracewell, R. N. (2000). The Fourier Transform And Its Applications. McGraw-Hill series in electrical and computer engineering. Circuits and systems. McGraw Hill, 3 edition.

Bruton, M. R., O’Dwyer, N., and Adams, R. (2013). Sex differences in the kinematics and neuromuscular control of landing: Biological, environmental and sociocultural factors. Journal of Electromyography and Kinesiology, 23(4):747–758. doi: 10.1016/j.jelekin.2013.04.012.

Corcoran, A. J., Schirmacher, M. R., Black, E., and Hedrick, T. L. (2021). ThruTracker: Open-Source Software for 2-D and 3-D Animal Video Tracking. bioRxiv. doi: 10.1101/2021.05.12.443854.

D’Eath, R. (2012). Repeated locomotion scoring of a sow herd to measure lameness: consistency over time, the effect of sow characteristics and inter-observer reliability. Animal Welfare, 21(2):219–231. doi: 10.7120/09627286.21.2.219.

D’Inca, R., Gras-Le Guen, C., Che, L., Sangild, P. T., and Le Huërou-Luron, I. (2011). Intrauterine growth restriction delays feeding-induced gut adaptation in term newborn pigs. Neonatology, 99(3):208–216. doi: 10.1159/000314919.

Edwards, S. and Baxter, E. (2015). Chapter 11. Piglet mortality: causes and prevention. In The gestating and lactating sow, pages 253–278.

Feldpausch, J. A., Jourquin, J., Bergstrom, J. R., Bargen, J. L., Bokenkroger, C. D., Davis, D. L., Gonzalez, J. M., Nelssen, J. L., Puls, C. L., Trout, W. E., and Ritter, M. J. (2019). Birth weight threshold for identifying piglets at risk for preweaning mortality. Translational Animal Science, 3(2):633–640. doi: 10.1093/tas/txz076.

Fernández-Carrión, E., Martínez-Avilés, M., Ivorra, B., Martínez-López, B., Ramos, A. M., and Sánchez-Vizcaíno, J. M. (2017). Motion-based video monitoring for early detection of livestock diseases: The case of African swine fever. PLOS ONE, 12(9):1–13. doi: 10.1371/journal.pone.0183793.

Figueiredo, J., Santos, C. P., and Moreno, J. C. (2018). Automatic recognition of gait patterns in human motor disorders using machine learning: A review. Medical Engineering & Physics, 53:1–12. doi: 10.1016/j.medengphy.2017.12.006.

Fourier, J. (1822). Theorie analytique de la chaleur, par M. Fourier. Chez Firmin Didot, père et fils.

Frick, R. W. (1996). The appropriate use of null hypothesis testing. Psychological Methods, 1(4):379. doi: 10.1037/1082-989X.1.4.379.

Ganley, K. J. and Powers, C. M. (2005). Gait kinematics and kinetics of 7-year-old children: a comparison to adults using age-specific anthropometric data. Gait & Posture, 21(2):141–145. doi: 10.1016/j.gaitpost.2004.01.007.

Gelman, A., Carlin, J., Stern, H., Dunson, D., Vehtari, A., and Rubin, D. (2020). Bayesian Data Analysis. 3rd edition. http://www.stat.columbia.edu/~gelman/book.

Gray, R. M. and Goodman, J. W. (1995). Fourier Transforms: An Introduction for Engineers. The Springer International Series in Engineering and Computer Science 322. Springer US, 1 edition.

Guesgen, M. and Bench, C. (2017). What can kinematics tell us about the affective states of animals? Animal Welfare, 26:383–397. doi: 10.7120/09627286.26.4.383.

Gómez, Y., Stygar, A. H., Boumans, I. J. M. M., Bokkers, E. A. M., Pedersen, L. J., Niemi, J. K., Pastell, M., Manteca, X., and Llonch, P. (2021). A systematic review on validated precision livestock farming technologies for pig production and its potential to assess animal welfare. Frontiers in Veterinary Science, 8. doi: 10.3389/fvets.2021.660565.

Hales, J., Moustsen, V. A., Nielsen, M. B. F., and Hansen, C. F. (2013). Individual physical characteristics of neonatal piglets affect preweaning survival of piglets born in a noncrated system. Journal of Animal Science, 91(10):4991–5003. doi: 10.2527/jas.2012-5740.

Hof, A. L. (1996). Scaling gait data to body size. Gait & Posture, 4(3):222–223. doi: 10.1016/0966-6362(95)01057-2.

Howson, E., Soldan, A., Webster, K., Beer, M., Zientara, S., Belak, S., Sanchez-Vizcaino, J., Van Borm, S., King, D., and Fowler, V. (2017).

Irschick, D. and Jayne, B. (1999). Comparative three-dimensional kinematics of the hindlimb for high-speed bipedal and quadrupedal locomotion of lizards. Journal of Experimental Biology, 202(9):1047–1065. doi: 10.1242/jeb.202.9.1047.

Jackson, B. E., Evangelista, D. J., Ray, D. D., and Hedrick, T. L. (2016). 3D for the people: multi-camera motion capture in the field with consumer-grade cameras and open source software. Biology Open, 5(9):1334–1342. doi: 10.1242/bio.018713.

Karashchuk, P., Rupp, K. L., Dickinson, E. S., Walling-Bell, S., Sanders, E., Azim, E., Brunton, B. W., and Tuthill, J. C. (2021). Anipose: A toolkit for robust markerless 3D pose estimation. Cell Reports, 36(13):109730. doi: 10.1016/j.celrep.2021.109730.

Lamy, E. and Mau, M. (2012). Saliva proteomics as an emerging, non-invasive tool to study livestock physiology, nutrition and diseases. Journal of Proteomics, 75(14):4251–4258. doi: 10.1016/j.jprot.2012.05.007. Special Issue: Farm Animal Proteomics.

LaVallee, K. T., Maus, T. P., Stock, J. D., Stalder, K. J., Karriker, L. A., Murthy, N. S., Kanwar, R., Beutler, A. S., and Unger, M. D. (2020). Quantitation of Gait and Stance Alterations Due to Monosodium Iodoacetate–induced Knee Osteoarthritis in Yucatan Swine. Comparative medicine, 70(3):248–257.

Le Dividich, J., Charneca, R., and Thomas, F. (2017). Relationship between birth order, birth weight, colostrum intake, acquisition of passive immunity and pre-weaning mortality of piglets. Spanish Journal of Agricultural Research, 15(2):e0603. doi: 10.5424/sjar/2017152-9921.

Litten, J., Drury, P., Corson, A., Lean, I., and Clarke, L. (2003). The influence of piglet birth weight on physical and behavioural development in early life. Neonatology, 84(4):311–318. doi: 10.1159/000073640.

Marchant, J. N., Rudd, A. R., Mendl, M. T., Broom, D. M., Meredith, M. J., Corning, S., and Simmins, P. H. (2000). Timing and causes of piglet mortality in alternative and conventional farrowing systems. Veterinary Record, 147(8):209–214. doi: 10.1136/vr.147.8.209.

Mathis, A., Mamidanna, P., Cury, K. M., Abe, T., Murthy, V. N., Mathis, M. W., and Bethge, M. (2018). DeepLabCut: markerless pose estimation of user-defined body parts with deep learning. Nature neuroscience, 21(9):1281–1289. doi: 10.1038/s41593-018-0209-y.

Mathis, A., Schneider, S., Lauer, J., and Mathis, M. W. (2020). A Primer on Motion Capture with Deep Learning: Principles, Pitfalls, and Perspectives. Neuron, 108(1):44–65. doi: 10.1016/j.neuron.2020.09.017.

Maxwell, S. E., Delaney, H. D., and Kelley, K. (2017). Designing Experiments and Analyzing Data: a Model Comparison Perspective. Routledge, New York.

McElreath, R. (2018). Statistical Rethinking: a Bayesian Course With Examples in R and Stan. Chapman and Hall/CRC.

Meijer, E., Bertholle, C. P., Oosterlinck, M., van der Staay, F. J., Back, W., and van Nes, A. (2014). Pressure mat analysis of the longitudinal development of pig locomotion in growing pigs after weaning. BMC veterinary research, 10(1):1–11. doi: 10.1186/1746-6148-10-37.

Michelini, A., Eshraghi, A., and Andrysek, J. (2020). Two-dimensional video gait analysis: A systematic review of reliability, validity, and best practice considerations. Prosthetics and Orthotics International, 44(4):245–262. doi: 10.1177/030936462092129.

Mielke, F., Schunke, V., Wölfer, J., and Nyakatura, J. A. (2018). Motion analysis of non-model organisms using a hierarchical model: Influence of setup enclosure dimensions on gait parameters of Swinhoe’s striped squirrels as a test case. Zoology, 129:35–44. doi: 10.1016/j.zool.2018.05.009.

Mielke, F., Van Ginneken, C., and Aerts, P. (2019). Quantifying intralimb coordination of terrestrial ungulates with Fourier coefficient affine superimposition. Zoological Journal of the Linnean Society, 189(3):1067–1083. doi: 10.1093/zoolinnean/zlz135.

Mielke, M., Aerts, P., Van Ginneken, C., Van Wassenbergh, S., and Mielke, F. (2020). Progressive tracking: a novel procedure to facilitate manual digitization of videos. Biology Open, 9(11). doi: 10.1242/bio.055962.

Mirkiani, S., Roszko, D. A., O’Sullivan, C. L., Faridi, P., Hu, D. S., Fang, D., Everaert, D. G., Toossi, A., Robinson, K., and Mushahwar, V. K. (2021). Overground Gait Kinematics and Muscle Activation Patterns in the Yucatan Mini Pig. bioRxiv. doi: 10.1101/2021.10.19.465020.

Muns, R., Manzanilla, E. G., Sol, C., Manteca, X., and Gasa, J. (2013). Piglet behavior as a measure of vitality and its influence on piglet survival and growth during lactation. Journal of Animal Science, 91(4):1838–1843. doi: 10.2527/jas.2012-5501.

Neethirajan, S. (2020). The role of sensors, big data and machine learning in modern animal farming. Sensing and Bio-Sensing Research, 29:100367. doi: 10.1016/j.sbsr.2020.100367.

Netukova, S., Duspivova, T., Tesar, J., Bejtic, M., Baxa, M., Ellederova, Z., Szabo, Z., and Krupicka, R. (2021). Instrumented pig gait analysis: State-of-the-art. Journal of Veterinary Behavior, 45:51–59. doi: 10.1016/j.jveb.2021.06.006.

Newell, K. M. and Liu, Y.-T. (2021). Collective Variables and Task Constraints in Movement Coordination, Control and Skill. Journal of Motor Behavior, 53(6):770–796. doi: 10.1080/00222895.2020.1835799.

Nishikawa, K., Biewener, A. A., Aerts, P., Ahn, A. N., Chiel, H. J., Daley, M. A., Daniel, T. L., Full, R. J., Hale, M. E., Hedrick, T. L., Lappin, A. K., Nichols, T. R., Quinn, R. D., Satterlie, R. A., and Szymik, B. (2007). Neuromechanics: an integrative approach for understanding motor control. Integrative and Comparative Biology, 47(1):16–54. doi: 10.1093/icb/icm024.

Patua, R., Muchhal, T., and Basu, S. (2021). Gait-Based Person Identification, Gender Classification, and Age Estimation: A Review. In Panigrahi, C. R., Pati, B., Mohapatra, P., Buyya, R., and Li, K.-C., editors, Progress in Advanced Computing and Intelligent Engineering, pages 62–74, Singapore. Springer Singapore.

Piñeiro, C., Morales, J., Rodríguez, M., Aparicio, M., Manzanilla, E. G., and Koketsu, Y. (2019). Big (pig) data and the internet of the swine things: a new paradigm in the industry. Animal Frontiers, 9(2):6–15. doi: 10.1093/af/vfz002.

Pike, A. V. L. and Alexander, R. M. (2002). The relationship between limb-segment proportions and joint kinematics for the hind limbs of quadrupedal mammals. Journal of Zoology, 258(4):427–433. doi: 10.1017/S0952836902001577.

Qiao, Y., Kong, H., Clark, C., Lomax, S., Su, D., Eiffert, S., and Sukkarieh, S. (2021). Intelligent Perception-Based Cattle Lameness Detection and Behaviour Recognition: A Review. Animals, 11(11). doi: 10.3390/ani11113033.

Quiniou, N., Dagorn, J., and Gaudré, D. (2002). Variation of piglets’ birth weight and consequences on subsequent performance. Livestock production science, 78(1):63–70. doi: 10.1016/S0301-6226(02)00181-1.

Riley, R. D., Ensor, J., Snell, K. I. E., Harrell, F. E., Martin, G. P., Reitsma, J. B., Moons, K. G. M., Collins, G., and van Smeden, M. (2020). Calculating the sample size required for developing a clinical prediction model. BMJ, 368. doi: 10.1136/bmj.m441.

Roehe, R. and Kalm, E. (2000). Estimation of genetic and environmental risk factors associated with pre-weaning mortality in piglets using generalized linear mixed models. Animal Science, 70(2):227–240. doi: 10.1017/S1357729800054692.

Salvatier, J., Wiecki, T. V., and Fonnesbeck, C. (2016). Probabilistic programming in Python using PyMC3. PeerJ Computer Science, 2:e55. doi: 10.7717/peerj-cs.55.

Schlageter-Tello, A., Bokkers, E. A., Groot Koerkamp, P. W., Van Hertem, T., Viazzi, S., Romanini, C. E., Halachmi, I., Bahr, C., Berckmans, D., and Lokhorst, K. (2014). Manual and automatic locomotion scoring systems in dairy cows: A review. Preventive Veterinary Medicine, 116(1):12–25. doi: 10.1016/j.prevetmed.2014.06.006.

Schmitt, O., Baxter, E. M., Lawlor, P. G., Boyle, L. A., and O’Driscoll, K. (2019). A Single Dose of Fat-Based Energy Supplement to Light Birth Weight Pigs Shortly After Birth Does Not Increase Their Survival and Growth. doi: 10.3390/ani9050227.

Scott, B., Seyres, M., Philp, F., Chadwick, E. K., and Blana, D. (2022). Healthcare applications of single camera markerless motion capture: a scoping review. doi: 10.7717/peerj.13517.

Seethapathi, N., Wang, S., Saluja, R., Blohm, G., and Kording, K. P. (2019). Movement science needs different pose tracking algorithms. arXiv, 1907.10226. doi: 10.48550/arXiv.1907.10226.

Serra Bragança, F., Rhodin, M., and van Weeren, P. (2018). On the brink of daily clinical application of objective gait analysis: What evidence do we have so far from studies using an induced lameness model? The Veterinary Journal, 234:11–23. doi: 10.1016/j.tvjl.2018.01.006.

Stavrakakis, S., Guy, J., Warlow, O., Johnson, G., and Edwards, S. (2014). Walking kinematics of growing pigs associated with differences in musculoskeletal conformation, subjective gait score and osteochondrosis. Livestock Science, 165:104–113. doi: 10.1016/j.livsci.2014.04.008.

Stiffler-Joachim, M. R., Wille, C., Kliethermes, S., and Heiderscheit, B. (2020). Factors Influencing Base of Gait During Running: Consideration of Sex, Speed, Kinematics, and Anthropometrics. Journal of Athletic Training, 55(12):1300–1306. doi: 10.4085/1062-6050-565-19.

van de Schoot, R., Depaoli, S., King, R., Kramer, B., Märtens, K., Tadesse, M. G., Vannucci, M., Gelman, A., Veen, D., Willemsen, J., et al. (2021). Bayesian Statistics and Modelling. Nature Reviews Methods Primers, 1(1):1–26. doi: 10.1038/s43586-020-00001-2.

Van Ginneken, C., Ayuso, M., Van Bockstal, L., and Van Cruchten, S. (2022). Preweaning performance in intrauterine growth-restricted piglets: Characteristics and interventions. Molecular Reproduction and Development. doi: 10.1002/mrd.23614.

Van Tichelen, K., Prims, S., Ayuso, M., Van Kerschaver, C., Vandaele, M., Degroote, J., Van Cruchten, S., Michiels, J., and Van Ginneken, C. (2021). Handling Associated with Drenching Does Not Impact Survival and General Health of Low Birth Weight Piglets. Animals, 11(2). doi: 10.3390/ani11020404.

Vanden Hole, C., Aerts, P., Prims, S., Ayuso, M., Van Cruchten, S., and Van Ginneken, C. (2018a). Does intrauterine crowding affect locomotor development? A comparative study of motor performance, neuromotor maturation and gait variability among piglets that differ in birth weight and vitality. PLOS ONE, 13(4):1–21. doi: 10.1371/journal.pone.0195961.

Vanden Hole, C., Ayuso, M., Aerts, P., Prims, S., Van Cruchten, S., and Van Ginneken, C. (2019). Glucose and glycogen levels in piglets that differ in birth weight and vitality. Heliyon, 5(9):e02510. doi: 10.1016/j.heliyon.2019.e02510.

Vanden Hole, C., Ayuso, M., Aerts, P., Van Cruchten, S., Thymann, T., Sangild, P. T., and Van Ginneken, C. (2021). Preterm Birth Affects Early Motor Development in Pigs. Frontiers in Pediatrics, 9:1003. doi: 10.3389/fped.2021.731877.

Vanden Hole, C., Cleuren, S., Van Ginneken, C., Prims, S., Ayuso, M., Van Cruchten, S., and Aerts, P. (2018b). How does intrauterine crowding affect locomotor performance in newborn pigs? A study of force generating capacity and muscle composition of the hind limb. PLOS ONE, 13(12):1–18. doi: 10.1371/journal.pone.0209233.

Vanden Hole, C., Goyens, J., Prims, S., Fransen, E., Ayuso Hernando, M., Van Cruchten, S., Aerts, P., and Van Ginneken, C. (2017). How innate is locomotion in precocial animals? A study on the early development of spatio-temporal gait variables and gait symmetry in piglets. Journal of Experimental Biology, 220(15):2706–2716. doi: 10.1242/jeb.157693.

von Wachenfelt, H., Pinzke, S., Nilsson, C., Olsson, O., and Ehlorsson, C.-J. (2008). Gait analysis of unprovoked pig gait on clean and fouled concrete surfaces. Biosystems Engineering, 101(3):376–382. doi: 10.1016/j.biosystemseng.2008.09.002.

Vranken, E. and Berckmans, D. (2017). Precision livestock farming for pigs. Animal Frontiers, 7(1):32–37. doi: 10.2527/af.2017.0106.

Wang, J., Yang, M., Cao, M., Lin, Y., Che, L., Duraipandiyan, V., Al-Dhabi, N. A., Fang, Z., Xu, S., Feng, B., Liu, G., and Wu, D. (2016). Moderately increased energy intake during gestation improves body condition of primiparous sows, piglet growth performance, and milk fat and protein output. Livestock Science, 194:23–30. doi: 10.1016/j.livsci.2016.09.012.

Wathes, C., Kristensen, H., Aerts, J.-M., and Berckmans, D. (2008). Is precision livestock farming an engineer’s daydream or nightmare, an animal’s friend or foe, and a farmer’s panacea or pitfall? Computers and Electronics in Agriculture, 64(1):2–10. doi: 10.1016/j.compag.2008.05.005. Smart Sensors in precision livestock farming.

Webb, D. and Sparrow, W. A. (2007). Description of joint movements in human and non-human primate locomotion using Fourier analysis. Primates, 48(4):277–292. doi: 10.1007/s10329-007-0043-4.

Wurtz, K., Camerlink, I., D’Eath, R. B., Fernández, A. P., Norton, T., Steibel, J., and Siegford, J. (2019). Recording behaviour of indoor-housed farm animals automatically using machine vision technology: A systematic review. PLOS ONE, 14(12):1–35. doi: 10.1371/journal.pone.0226669.

Yitbarek, D. and Dagnaw, G. G. (2022). Application of Advanced Imaging Modalities in Veterinary Medicine: A Review. Veterinary Medicine: Research and Reports, 13:117. doi: 10.2147/VMRR.S367040.

Young, J. W. and Shapiro, L. J. (2018). Developments in development: What have we learned from primate locomotor ontogeny? American Journal of Physical Anthropology, 165(S65):37–71. doi: 10.1002/ajpa.23388.

